# 3Dmapper: A Command Line Tool For BioBank-scale Mapping Of Variants To Protein Structures

**DOI:** 10.1101/2023.09.01.555502

**Authors:** Victoria Ruiz-Serra, Samuel Valentini, Sergi Madroñero, Alfonso Valencia, Eduard Porta-Pardo

**Affiliations:** Barcelona Supercomputing Center (BSC); Josep Carreras Leukaemia Research Institute (IJC), Badalona, Spain; Department of Cellular, Computational and Integrative Biology (CIBIO), University of Trento, Trento, Italy; Institució Catalana de Recerca Avançada (ICREA)

**Keywords:** Variant mapping, Protein structures, Command line tool, Genomic data interpretation, Gene-to-protein mapping

## Abstract

The interpretation of genomic data is crucial to understand the molecular mechanisms of biological processes. Protein structures play a vital role in facilitating this interpretation by providing functional context to genetic coding variants. However, mapping genes to proteins is a tedious and error-prone task due to inconsistencies in data formats. Over the past two decades, numerous tools and databases have been developed to automatically map annotated positions and variants to protein structures. However, most of these tools are web-based and not well-suited for large-scale genomic data analysis. To address this issue, we introduce 3Dmapper, a standalone command-line tool developed in Python and R. It systematically maps annotated protein positions and variants to protein structures, providing a solution that is both efficient and reliable. 3Dmapper is freely available on GitHub at https://github.com/vicruiser/3Dmapper.

## SUMMARY

Mutations in protein coding regions are particularly significant as they can cause functional damage to proteins. This is clearly evidenced by the fact that most Mendelian disease and the vast majority of highly penetrant disease-associated variants are located in protein coding regions^1^. However, while we have made significant progress in predicting the consequences of many types of different mutations, understanding the degree of damage of coding genetic variants remains a particularly challenging task.

To that end, there are now dozens of different tools that aim to help researchers to either predict or understand the consequences of such genetic variants. Some of these rely solely on the linear sequence information of the protein and the variant^2–4^. While these have the advantage of being able to be applied to virtually any genetic variant, the accuracy and precision of their output tends to be lower compared to tools that exploit the three-dimensional structure of the proteins^5^. The reason is that the structural context of genetic variants provides more information about the potential function of the affected residues and the likely consequences of the variants. This has been proven key to, among others, distinguish between driver and passenger mutations in cancer genes^5^ or identify germline pathogenic variants^5^.

One significant limitation of methods that exploit three-dimensional structure, however, is that we currently only have experimental structures for a subset of the human proteome^6^. Furthermore, the mapping between genomic coordinates and three-dimensional structures is not trivial. For example, oftentimes there are mismatches between amino acid positions in Protein Data Bank (PDB)^7^ structure files and those in protein sequences from databases like Uniprot^11^or Ensembl^9^. Furthermore, discrepancies can arise when the protein used for the structural studies does not match the reference protein in the databases or in the genome sequences due to mutations introduced to facilitate the structural studies.

Over the past two decades, there has been significant progress in developing tools and databases to automatically map annotated positions or variants to protein structures (see **Table S1**). However, many of these tools suffer from limitations including being outdated or not actively maintained. Furthermore, keeping up with data updates, including new variants and the latest protein structures, can be a challenge. These limitations hinder the widespread application of these methods in the current era of large-scale genomics projects, where researchers routinely face the analysis of thousands of genomes or exomes at once.

Fortunately, there is ongoing development and periodic release of new tools to address these issues. However, it is important to note that some tools are focused solely on human data, while others only map mutations without considering genomic or protein positions of interest in the broader proteome or other organisms’ proteomes. Additionally, many of the available tools are web-based, which is convenient for non-expert users lacking programming skills but may not be suitable for large-scale genomic data analysis.

Here we introduce 3Dmapper, a versatile standalone command-line tool in Python and R that maps annotated protein positions or variants to protein structures. 3Dmapper accepts PDB format files (.pdb and .cif) for any organism and allows users to control the data version. Our tool implements a homology sequence search approach to find structural templates when a protein structure is unavailable. It also provides functional annotation of protein-protein, protein-ligand, and protein-nucleic interfaces by determining inter-residue distances, facilitating the functional interpretation of mapped positions. Additionally, 3Dmapper includes associated B-factors for each mapped residue, corresponding to the pLDDT model quality metric in AlphaFold2 models^10^, providing valuable information should the user want to exploit AlphaFold2 models.

## METHODS

### Installation and dependencies

To download and install 3Dmapper, please visit the following link: https://github.com/vicruiser/3Dmapper. A detailed tutorial on how to use the tool with examples is available at the same link as well as in the **Supplementary Information**. 3Dmapper depends on R≥3.5, Python≥3.6, and BLAST>=2.6.

### 3Dmapper pipeline

The 3Dmapper pipeline (**Figure 1**) consists of four command-line tools:

- *makestructuraldb*: it generates a structurally annotated protein database by aligning user-defined protein sequences and structures with BLAST. It offers the option to
- customize the sequence identity cutoff (Pident) to maximize the inclusion of structural homologs. Then, it calculates inter-chain interfaces based on user-defined spatial proximity and generates individual output files with detailed structural information per protein ID.
- *makevariantsdb*: takes variants or annotated positions files in .vcf, .vep or .maf format and simply splits them by transcript ID. This process is useful for speeding up the subsequent mapping process.
- *mapper*: maps variants or protein positions to protein structures using the precomputed structural and variants files from *makestructuraldb* and *makevariantsdb* respectively. It generates a table in .csv or .hdf5 along with a SetID file, being the latter useful for rare-variant association testing^11^
- *makevisualization*: generates ChimeraX^12^ scripts for automatic visualizations of the mapped variants/positions using *mapper*’s output.

**Figure 1.**
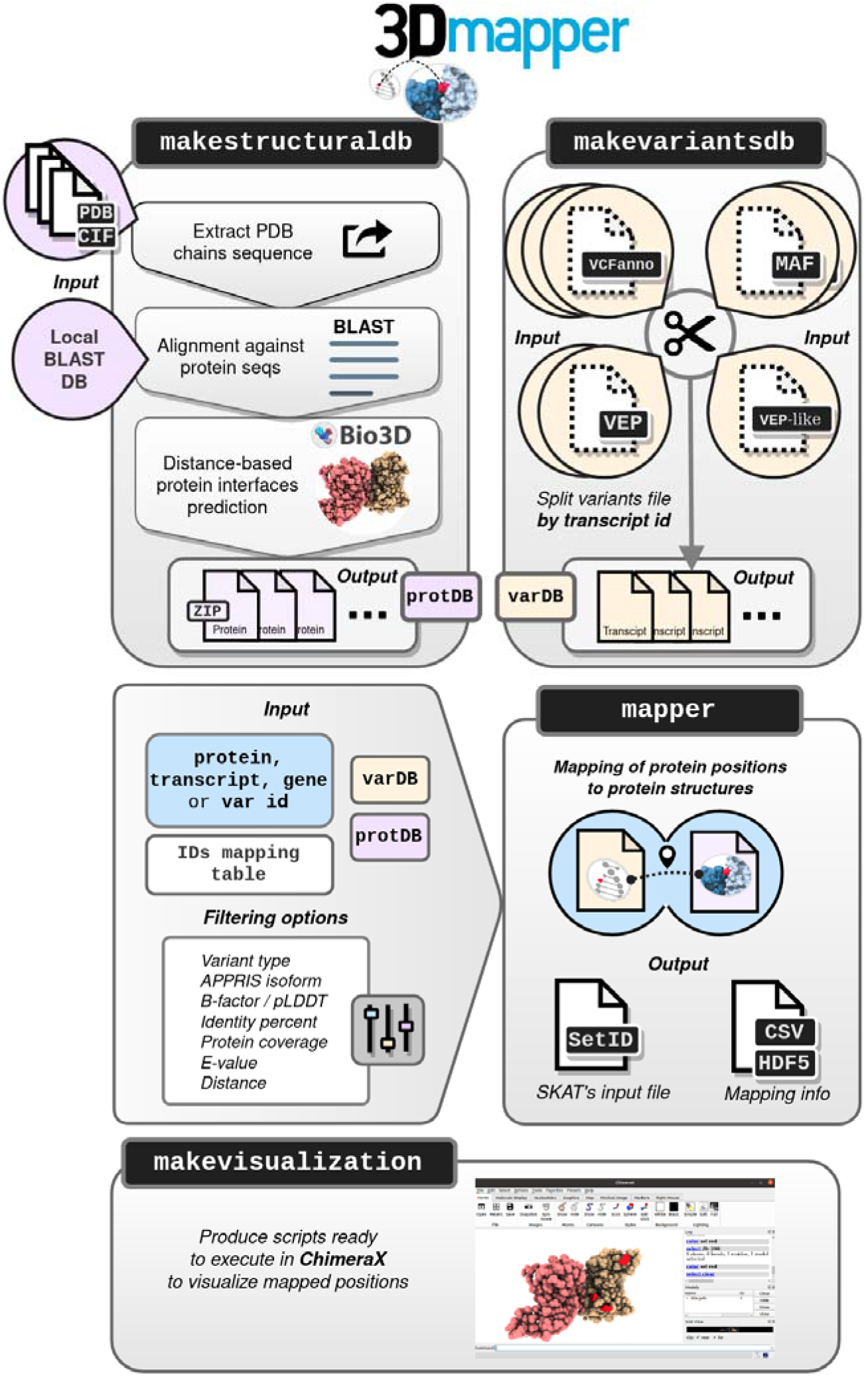
The workflow of 3Dmapper. The pipeline begins with the generation of a structural dataset using *makestructuraldb*, which is then utilized to map genetic variants or protein positions using *mapper*. To facilitate the mapping process, variants are first split into smaller files using *makevariantsdb*. Additionally, ChimeraX visualization scripts of the mapped positions can be generated automatically using *makevisualization*. VCFanno: VCF type of annotation; MAF: Mutation Annotation Format; VEP: Variant Effect Predictor output file; VEP-like: similar to VEP output but with extra or missing columns compared to the original one; varDB: variants database; protDB: protein database.

More details about how to run each of the commands, input and output formats and examples are provided in **Supplementary Information** and in the GitHub wiki.

### 3Dmapper performance

We evaluated the performance of 3Dmapper to analyze its time and memory usage, using Python’s ‘memory profiler’ module (https://pypi.org/project/memory-profiler/). This evaluation offers insights into the efficiency and resource requirements of the mapping workflow process.

### Structural database

A comprehensive human protein structural database was generated using *makestructuraldb*. This database incorporated the human proteome from Uniprot (https://www.uniprot.org/proteomes/UP000005640) and the full PDB, comprising 194,551 files in .cif format as of 30/08/2022. We used default input parameters and set Pident to 50% and e-value cutoff of ≤10^-5^ to ensure reliable structural templates for human proteins.

### Coding genetic variants

To map to the human protein structural database, variants were sourced from publicly available repositories, including TCGA (https://api.gdc.cancer.gov/data/1c8cfe5f-e52d-41ba-94da-f15ea1337efc), gnomAD v2.1.1^13^ (https://storage.googleapis.com/gcp-public-data--gnomad/release/2.1.1/vcf/exomes/gnomad.exomes.r2.1.1.sites.vcf.bgz), and ClinVar^14^ (https://ftp.ncbi.nlm.nih.gov/pub/clinvar/vcf_GRCh38/clinvar.vcf.gz). Additionally, the UK BioBank 50,000 whole-exomes dataset from March 2019 (UKBB) was included. When protein locations of variants were not available, Ensembl Variant Effect Predictor (VEP; version 98.3) was used to predict them, particularly for variants in .vcf format. Clinical interpretation data for ClinVar and TCGA variants were sourced from https://ftp.ncbi.nlm.nih.gov/pub/clinvar/tab_delimited/variant_summary.txt.gz and the referenced file, respectively.

### Comparison between 3Dmapper with interactome3D interfaces

We used the human Interactome3D dataset^15^ v2020_05 to benchmark the interfaces predicted by 3Dmapper.

### Removal of protein interface redundancy

To eliminate protein interfaces’ redundancy, we filter out interfaces that exhibit less than 80% Pident between the PDB and the Uniprot sequences. When multiple PDB structures exist for a given protein, we prioritize the PDB structure with the highest Pident for each type of interface, including protein, ligand, and nucleic interfaces. This approach enables us to capture interface regions spanning the entire protein. For ligands, we treat each individual ligand as a distinct interface.

### AlphaFold models of human protein complexes

We obtained predicted protein-protein complexes from the hu.MAP^16^ and HuRI^17^ databases, as described in a recent publication^18^. The data were downloaded from https://archive.bioinfo.se/huintaf2/ and processed using the *makestructuraldb* command to create a structurally annotated protein database (available at Zenodo^19^). Subsequently, variants were mapped to the AlphaFold complexes using the *mapper* command. We considered the available pDockQ scores as a proxy for the quality of the models, as they take into account both pLDDT and inter-residue distance^20^. Thus, residues with pDockQ scores greater than 0.23, indicating acceptable models based on experimental validation, were included in the analysis^18^.

### Software

Graphical plots were done using R 3.6.3 with the package ggplot2^21^. All the PDBs handling in *makestructuraldb* is done with the Bio3D R package^22^. Parallelization of jobs in the computer cluster MareNostrum4 were managed with Greasy 2.2.4 (https://github.com/jonarbo/GREASY).

## RESULTS

To evaluate the efficiency and performance of 3Dmapper, we conducted comprehensive tests to assess its ability to handle large numbers of variants (ClinVar, gnomAD, TCGA and 50.000 whole-exomes from the UK BioBank) and map them to the full PDB database (see Methods). This evaluation not only provides insights into the time and memory requirements of the tool but also serves as a practical guide to map a biobank-scale set of variants to PDB structures.

We first used the 3Dmapper’s *makestructuraldb* command to generate a structural database for the human proteome in Uniprot and related PDB files. We executed the process using 672 CPUs in parallel (Figure S1). The task was completed within 13 hours, resulting in a 13GB structural database accessible at Zenodo^19^. Results of makestructuraldb performance can be found in **Figure S1** and Methods. **Figure S2** shows an overview of the database content in terms of structural coverage for residues, proteins in PDBs and the significance of considering different levels of percent identity (Pident) for comprehensive protein structure homology analysis. Overall, approximately 10% of all residues have structural coverage, which increases by around 30% when considering 50% Pident. At Pident = 100%, around 35% of human proteins have structural coverage, increasing to nearly 70% at Pident = 50%.

To evaluate the quality interfaces calculated by 3Dmapper, we compared the predicted protein interactions from *makestructuraldb* with those from interactome3D^15^ (see Methods), a database integrating data from nine major public protein-interaction databases. We were able to identify 92% of the interactions described in interactome3D (n=10,361, **Figure S3**), along with over 350,000 additional protein-protein interactions based on distance, increasing the structural coverage of PPIs more than 30 times.

To explore how different types of genetic variants are distributed across the different structural features, we used the *makevariantsdb* command to split the VEP files containing variants from the four selected datasets. We conducted repeated tests for each variant database (TCGA, ClinVar, gnomAD, and UKBB) using different node configurations: 4, 8, and 16 CPUs. Each configuration was executed 10 times, with varying processing times proportional to the number of variants in the input file (**Figure S4**).

Next, we mapped the variants to the previously generated structural database using the *mapper* command. Considering one isoform per gene of Uniprot IDs that could be mapped to Ensembl transcript IDs, a total of 19,263 transcript IDs were processed. The command *mappe*r was executed in parallel with the option enabled for categorizing variants into interface, structure, unmapped, and non-coding classes and without filtering per type of variant. It utilized 96 CPUs in parallel, with 2 CPUs assigned to each task and 2GB of available RAM memory per CPU. The entire process took approximately 13 hours, with specific processing times for ClinVar (∼2 hours and 30 minutes), gnomAD (6 hours and 15 minutes), TCGA (∼2 hours), and UKBB (∼2 hours and 30 minutes) (**Figure S5**). The output sizes, considering interface and structure variants only, were ClinVar 14.8GB, gnomAD 220GB, TCGA 58GB, and UKBB 44GB. The computational times mentioned, along with the resulting file sizes, are reasonable and aligned with the demands of handling large-scale genetic data analysis.

Quantitative assessment of structural coverage based on Pident revealed that ClinVar, TCGA, and UKBB datasets had greater coverage compared to gnomAD when compared to the human proteome (**Figure S6A**). Disease-associated variants are strongly enriched in protein interfaces, when compared to non-disease associated variants (∼10% vs 1%) in terms of interfaces (**Figure S6B**). This is consistent with previous studies showing a strong enrichment of missense pathogenic variants at protein interfaces^23^.

Next, we compared mapping results across the four selected studies, focusing on missense variant coverage and their mapping to the structure or interfaces. Despite differences in the total number of variants considered, protein coverage was comparable in TCGA, UKBB, and gnomAD (**Figure S7A**).Integrating AlphaFold protein complex models of acceptable and high quality (pDockQ>0.23, see Methods) resulted in an approximate 30% increase in missense variants mapping for structure and interfaces in TCGA, UKBB, and gnomAD, and a 16% increase for Clinvar (Figure S7B). We attribute this effect to the total number of variants considered, the type of variants studied (clinical relevance), and the protein complexes considered in HuMap and HuRI.

As an example of how AlphaFold models and 3Dmapper can be useful, we studied mutations in the DNA repair protein XRCC2, which interacts with RAD51D to repair DNA double-strand breaks^24^ and is associated with cancer-related mutations (https://www.mycancergenome.org/content/gene/xrcc2/). This interaction lacked experimental structure or homology models in the considered PDB version. Among the ClinVar variants mapped to XRCC2, 90% were classified as VUS, with 16% of these variants localized within the protein interface (**Figure S8A**), providing improved interpretation possibilities. Additionally, TCGA variants like H86N and E97Q within the complex interface were predicted to have a damaging effect (**Figure S8B**). These alterations, involving a change from a basic to a polar amino acid at position 86 and from an acidic to a polar amino acid at position 97, could have the potential to disrupt the proper protein interaction, highlighting a possible functional significance. Other analyses highlighting the potential of 3Dmapper in different contexts can be found in recent publications, such as the analysis of somatic mutations in interfaces and their effects in disrupting protein protein interactions using data from the Clinical Proteomics Tumor Analysis Consortium (CPTAC) ^25^, or the analysis of evolutionary relevant positions across protein families mapped onto reference structures to analyze their local energetics, known as “frustration”, adding a layer of understanding to protein evolution dynamics ^26^.

## CONCLUSIONS

In summary, our software 3Dmapper solves the problem of mapping nonsense and missense mutations from the sequences to the protein structures, facilitating the systematic analysis of mutations in genomes, i.e. cancer genomes and providing a solid entry point to the prediction of the consequences of mutations. To demonstrate/validate the capacity of 3Dmapper, we have used it to map all the variants in the TCGA, ClinVar, UKBB, and gnomAD databases, to the corresponding structures of AlphaFold models, showing a between 16 to 30% improvement of the mapping percentages, especially for gnomAD variants.

3Dmapper represents a starting point for the systematic investigation of large-scale genomic and proteomic data, facilitating the study of the impact of mutations, a significant problem, since most Mendelian disease, and the vast majority of highly penetrant disease-associated variants, are located in protein coding regions^1^.

## Supporting information

3D mapper - Tutorial

## ACKNOWLEDGEMENTS

E.P-P and V.R-S are supported by the La Caixa Junior Leader Fellowship LCF/BQ/PI18/11630003. E.P-P also received support from the Spanish Ministry of Science (RYC2019-026415-I and PID2019-107043RA-I00). A.V. is supported by Institució Catalana de Recerca Avançada (ICREA). S.M. received support from the Asociación Española Contra el Cáncer (AECC) project LABAE20038PORT. The authors acknowledge the use of the Mare Nostrum 4 supercomputer cluster at the Barcelona Supercomputing Center (BSC) for the computational resources provided to carry out this research. This research has been conducted using the UK Biobank Resource under Application Number 54343. The results published here are in part based upon data generated by the TCGA Research Network (https://www.cancer.gov/tcga).

## REFERENCES

1. Bamshad, M. J. et al. Exome sequencing as a tool for Mendelian disease gene discovery. Nat. Rev. Genet. 12, 745–755 (2011).

2. Yang, F. et al. Protein domain-level landscape of cancer-type-specific somatic mutations. PLoS Comput. Biol. 11, e1004147 (2015).

3. Porta-Pardo, E. & Godzik, A. Mutation drivers of immunological responses to cancer. Cancer Immunol. Res. 4, 789–798 (2016).

4. Peterson, T. A. et al. DMDM: domain mapping of disease mutations. Bioinforma. Oxf. Engl. 26, 2458–2459 (2010).

5. Bailey, M. H. et al. Comprehensive Characterization of Cancer Driver Genes and Mutations. Cell 173, 371–385.e18 (2018).

6. Porta-Pardo, E., Ruiz-Serra, V., Valentini, S. & Valencia, A. The structural coverage of the human proteome before and after AlphaFold. PLOS Comput. Biol. 18, e1009818 (2022).

7. Berman, H. M. et al. The Protein Data Bank. Nucleic Acids Res. 28, 235–242 (2000).

8. The UniProt Consortium. UniProt: the Universal Protein Knowledgebase in 2023. Nucleic Acids Res. 51, D523–D531 (2023).

9. Hubbard, T. et al. The Ensembl genome database project. Nucleic Acids Res. 30, 38–41 (2002).

10. Jumper, J. et al. Highly accurate protein structure prediction with AlphaFold. Nature 596, 583–589 (2021).

11. Wu, M. C. et al. Rare-Variant Association Testing for Sequencing Data with the Sequence Kernel Association Test. Am. J. Hum. Genet. 89, 82–93 (2011).

12. Goddard, T. D. et al. UCSF ChimeraX: Meeting modern challenges in visualization and analysis. Protein Sci. Publ. Protein Soc. 27, 14–25 (2018).

13. Karczewski, K. J. et al. The mutational constraint spectrum quantified from variation in 141,456 humans. Nature 581, 434–443 (2020).

14. Landrum, M. J. et al. ClinVar: improving access to variant interpretations and supporting evidence. Nucleic Acids Res. 46, D1062–D1067 (2018).

15. Mosca, R., Céol, A. & Aloy, P. Interactome3D: adding structural details to protein networks. Nat. Methods 10, 47–53 (2013).

16. Drew, K., Wallingford, J. B. & Marcotte, E. M. hu.MAP 2.0: integration of over 15,000 proteomic experiments builds a global compendium of human multiprotein assemblies. Mol. Syst. Biol. 17, e10016 (2021).

17. Luck, K. et al. A reference map of the human binary protein interactome. Nature 580, 402–408 (2020).

18. Burke, D. F. et al. Towards a structurally resolved human protein interaction network. Nat. Struct. Mol. Biol. 30, 216–225 (2023).

19. Ruiz, V. 3Dmapper structural datasets. (2023) doi:10.5281/zenodo.8301664.

20. Bryant, P., Pozzati, G. & Elofsson, A. Improved prediction of protein-protein interactions using AlphaFold2. Nat. Commun. 13, 1265 (2022).

21. Wickham, H. Introduction. in ggplot2: Elegant Graphics for Data Analysis (ed. Wickham, H.) 1–7 (Springer, 2009). doi:10.1007/978-0-387-98141-3_1.

22. Grant, B. J., Skjærven, L. & Yao, X. The Bio3D packages for structural bioinformatics. Protein Sci. Publ. Protein Soc. 30, 20–30 (2021).

23. Cheng, F. et al. Comprehensive characterization of protein–protein interactions perturbed by disease mutations. Nat. Genet. 53, 342–353 (2021).

24. Baldock, R. A. et al. RAD51D splice variants and cancer-associated mutations reveal XRCC2 interaction to be critical for homologous recombination. DNA Repair 76, 99–107 (2019).

25. Li, Y. et al. Pan-cancer proteogenomics connects oncogenic drivers to functional states. Cell S0092–8674(23)00780–8 (2023) doi:10.1016/j.cell.2023.07.014.

26. Freiberger, M. I. et al. The Evolution of Local Energetic Frustration in Protein Families. 2023.01.25.525527 Preprint at 10.1101/2023.01.25.525527 (2023).

