## Supplementary material for "3Dmapper: A Command Line Tool For BioBank-scale Mapping Of Variants To Protein Structures": Suplementary Figures and Tables

**Table S1. Comparison of methods for mapping annotated variants or protein positions to protein structures.** Green and red backgrounds indicate whether the software is currently available and maintained or not, respectively. An orange background indicates that the website is available but results could not be obtained at the time of testing due to malfunctions. In the last column, the "O" designation signifies a specific organism of focus, while "A" represents any organism, "S" indicates selected organisms, and "H" designates exclusively human data. This table highlights the current status of each method and can serve as a guide for researchers in selecting the appropriate tool for their specific needs. Notably, within the "*Mapping protein positions & annotated mutations*" category, 3Dmapper stands as the only command line tool, underscoring the scarcity of tools of this type.

| **Method type** | **Tool name** | **Year** | **Description** | **O** |
| --- | --- | --- | --- | --- |
| *Mapping of protein positions* | | | | |
| **Web-based** | Seq2Struct^1^ | 2005 | Cross-link between PDB or SCOP with SwissProt and TrEMBL | A |
|  | PDBSWS^2^ | 2005 | Residue level mapping between PDB and Uniprot or SwissProt | A |
|  | FeatureMap3D^3^ | 2006 | Visualization of sequence features onto the structure of the best PDB hit | A |
|  | SSMap^4^ | 2008 | Residue level mapping between PDB and Uniprot | A |
|  | ProSAT+^5^ | 2016 | Interactive visualization of sequence positions to protein structures | A |
|  | G2S^6^ | 2018 | Residue level mapping between PDB and protein sequences | A |
|  | PDBrenum^7^ | 2021 | Renumbers residues in PDB files to match Uniprot residue numbering. Also in the format of a command line tool. | A |
| **Database** | SIFTS^8^ | 2012 | Residue level mapping between PDB and Uniprot | A |
| *Mapping of annotated mutations* | | | |  |
| **Web-based** | LS-SNP/PDB^9^ | 2009 | Mapping of human non-synonymous SNPs to PDB structures | H |
|  | CRAVAT^10^ | 2013 | Projection of cancer somatic mutations on available structures and homology models | H |
|  | SNP2Structure^11^ | 2015 | Map missense mutations to protein structures | H |
|  | G23D^12^ | 2016 | Map human genomic variants to protein structures | H |
|  | StructMAn^13^ | 2016 | Map SNVs to protein structures including possible homologs | A |
|  | COSMIC-3D^14^ | 2017 | Visualize COSMIC cancer mutations on 3D protein structures | H |
|  | GenProBis^15^ | 2017 | Interactive visualization of variants and genomic positions to individual protein structures. Provides information about ligand and protein binding sites. | S |
|  | VarQ^16^ | 2018 | Maps and predict the effect of Clinvar and Uniprot variants on protein structures | H |
|  | VarMap^17^ | 2019 | Map human genomic coordinates to protein sequence and structures | H |
|  | VarSite^18^ | 2020 | Detailed visualization of human disease-associated variants on PDBs | H |
|  | Venus^19^ | 2022 | Predicts the effects of non-synonymous coding variants on protein stability, post-translational modifications, linear motifs, and nearby residues | A |
|  | Swiss-PO^20^ | 2021 | Map human oncogenic mutations to protein structures | H |
| **Database** | MutDB^21^ | 2003 | 8000 human disease-associated SNPs from UCSC with structural annotation | H |
|  | MSV3d^22^ | 2012 | Human missense mutations mapped to 3D structures | H |
|  | cBioPortal^23^ | 2012 | Provides an interpretation of mutations using protein annotations and offers the choice of 3D visualization. | H |
|  | dSysMap^24^ | 2015 | Human disease-associated missense mutations mapped to Interactome3D with possibility of visualizing on corresponding protein structure interactively | H |
|  | Cancer3D^25^ | 2015 | Cancer associated mutations mapped to PDB structures | H |
|  | ADDRESS^26^ | 2021 | Human disease-associated and benign mutations from HUMSAVAR mapped to PDB structures | H |
| **Command line** | PDBMap^27^ | 2020 | Map genomic human variants onto protein structures | H |
| *Mapping protein positions & annotated mutations* | | | |  |
| **Web-based** | MuPIT^28^ | 2013 | Interactive visualization of variants and genomic positions to protein structures | H |
|  | ZoomVar^29^ | 2021 | Map SNVs and positions to human protein structures | H |
| **Stand-alone** | Jalview^30^ | 2009 | Connects coding DNA sequences within exons to protein sequences and structure | A |
| **Command line** | **3Dmapper** | 2023 | Tool to systematically map annotated genomic positions to protein structures in a large scale | A |

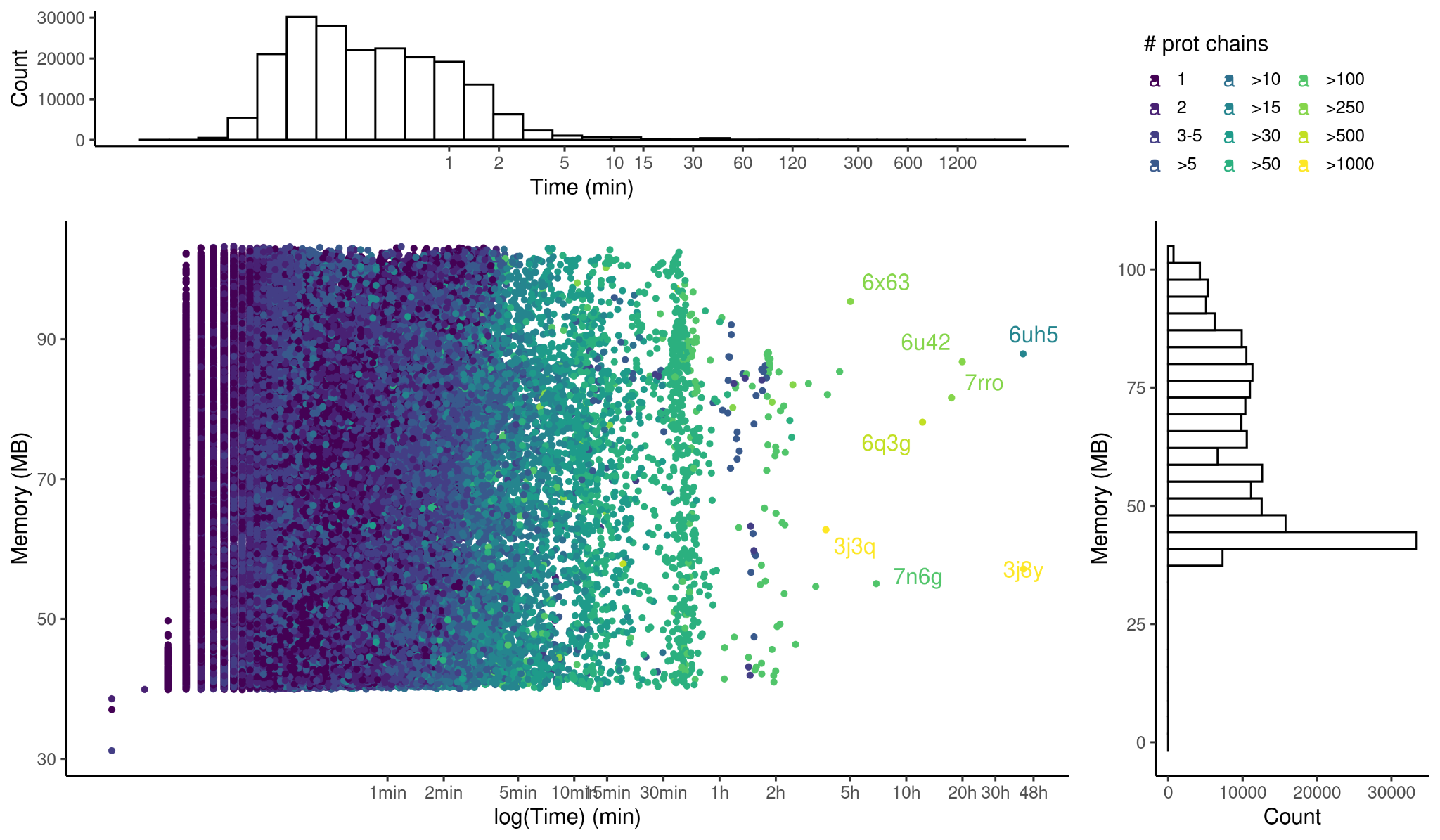

**Figure S1. Time and memory performance of *makestructuraldb*.** Each dot represents the processing of all chains of individual PDB files, and the color indicates the number of protein chains per PDB. Usually, more chains correspond to higher computing times, but the time is also affected by the length of the protein. The x-axis is in log2 scale. Most PDBs are processed within 5 minutes, except for a few outliers that take longer due to their size. These include PDBs with a high number of protein chains, such as 3J3Q (1356 chains) and 7RRO (429 chains), as well as those with very long proteins, such as 7N6G (3192 amino acids is the mean length of the 244 chains). The size of each individual processing does not exceed 100MB, and the majority are below 50MB, which is a reasonable use of memory for both, a cluster and a desktop computer.

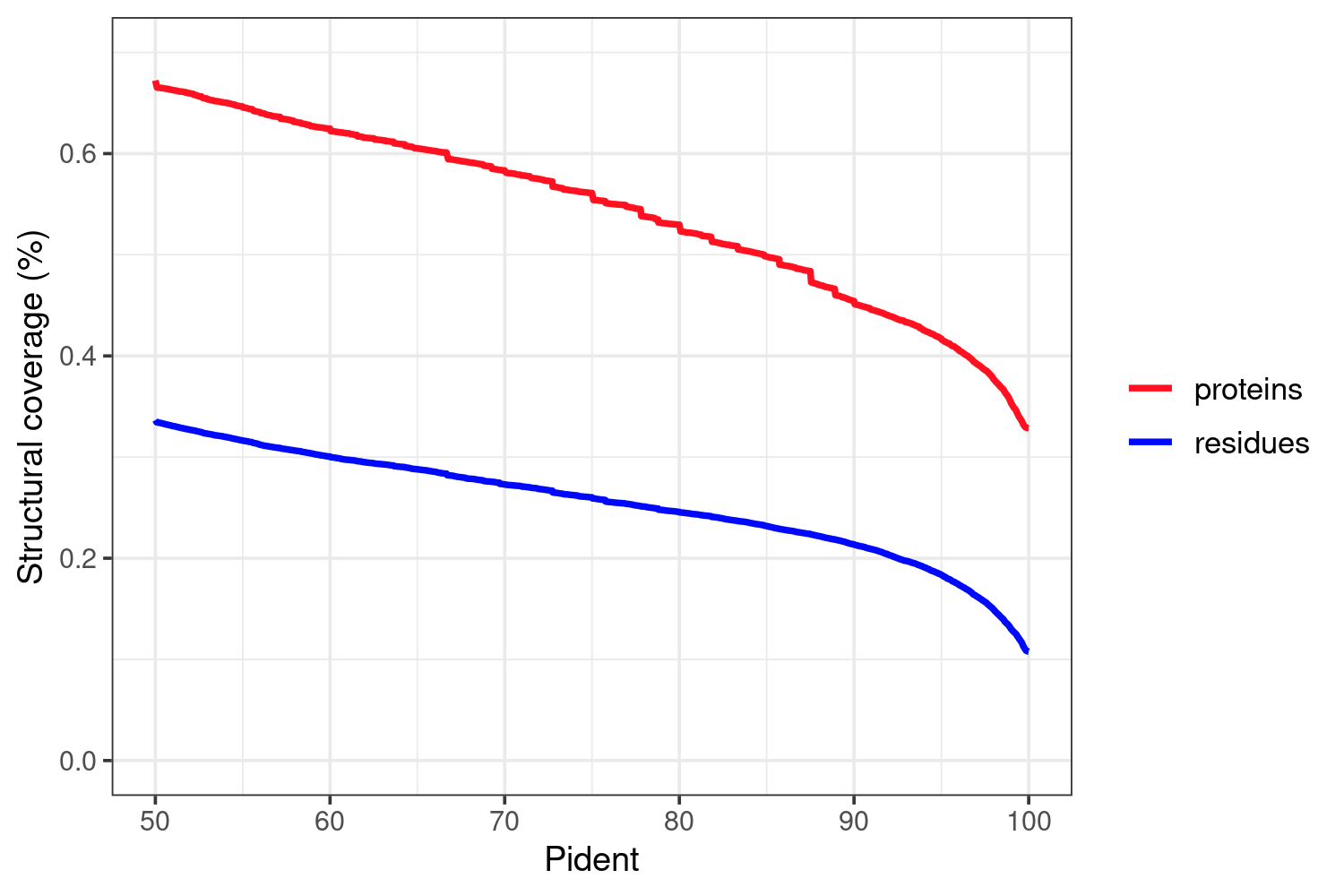

**Figure S2. Relationship between percent identity (Pident) and the structural coverage of the human proteome.** Human structural coverage at the residue level (n= 11395157) and protein level (one protein sequence per gene, n= 20600).

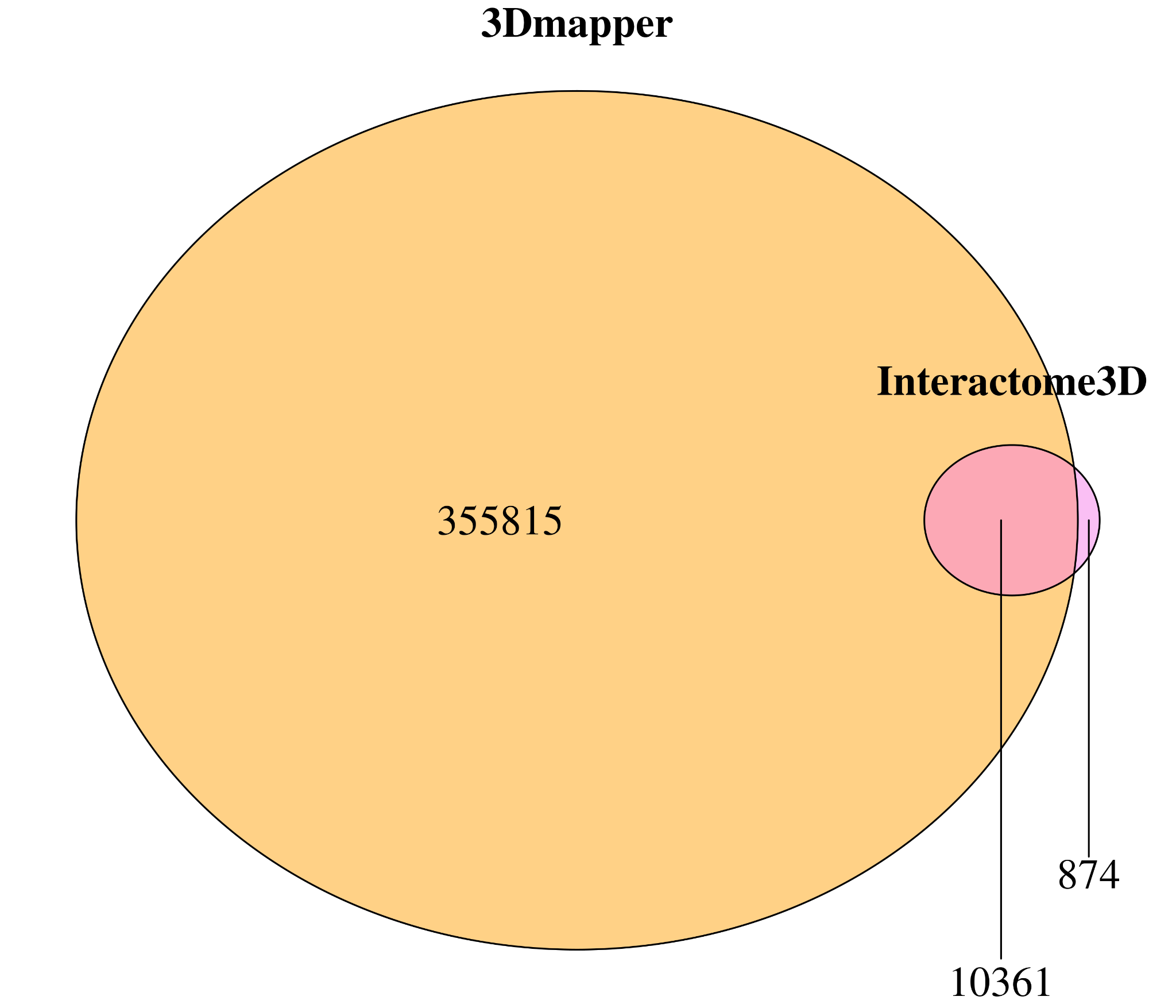

**Figure S3.** Venn Diagram showing the comparison of the protein-protein interactions (PPIs) predicted by Human Interactome3D and 3Dmapper, with a Pident threshold of 50%.

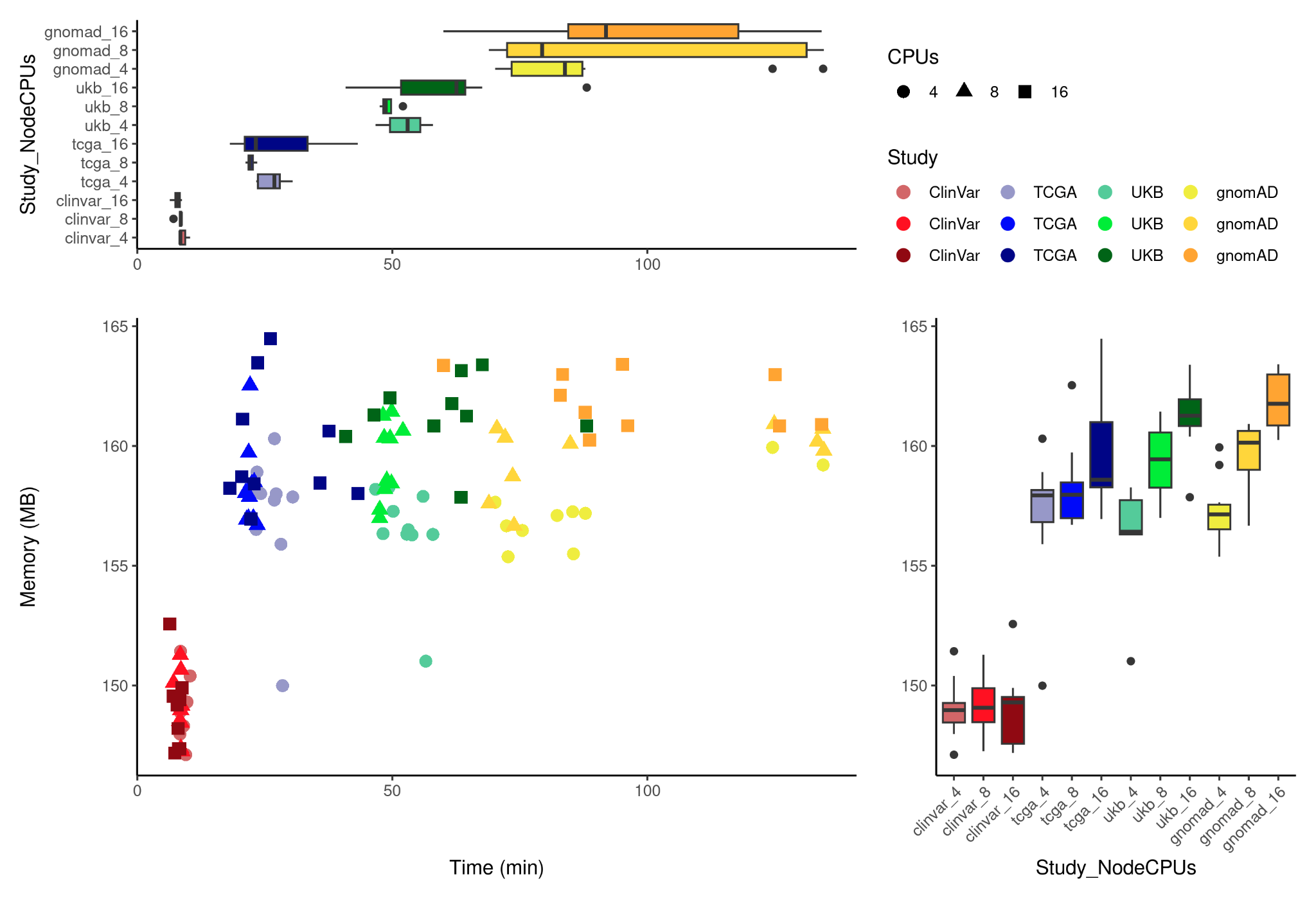

**Figure S4.** Time and memory performance of *makevariantsdb*, as assessed by executing the tool 10 times using 4, 8, and 16 CPUs on each of the four studies (Clinvar, TCGA, UKBB, gnomAD). The scatter plot shows the distribution of results, with the shape and color of the dots representing the number of CPUs and study, respectively. The accompanying boxplots summarize the time (top panel) and memory (left panel) distribution of the results. These findings provide insight into the optimal configuration for running *makevariantsdb* and can inform researchers' decisions when using this tool. despite the substantial number of coding variants (7,685,143 in gnomAD, 6,955,961 in UKBB, 2,344,782 in TCGA, and 596,161 in ClinVar), none of the executions took longer than 3 hours, indicating that the processing time remained reasonable even when dealing with a large number of variants. The computing time was found to be proportional to the number of variants, while the memory consumption remained relatively low (lower than 165MB).

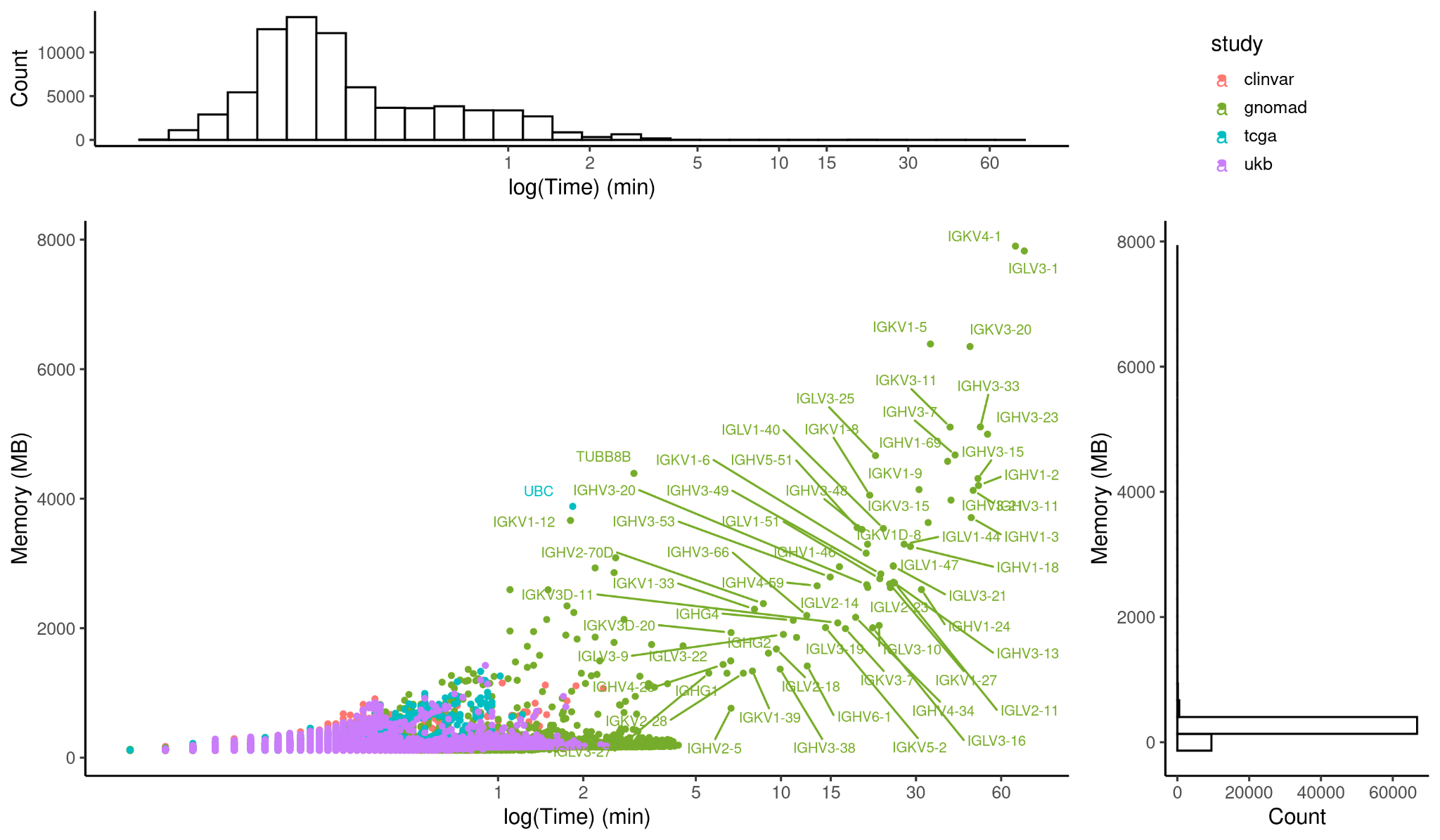

**Figure S5.** Time and memory performance of *mapper*. Each dot represents the mapping of variants from four different studies (color coded) to a total of 19,263 individual transcript IDs. The x-axis represents the time performance in log2 scale, while the y-axis represents the memory usage in MB. Labels of genes related to each transcript are depicted for jobs that took longer than 5 min or consumed more than 3.5GB of RAM memory.

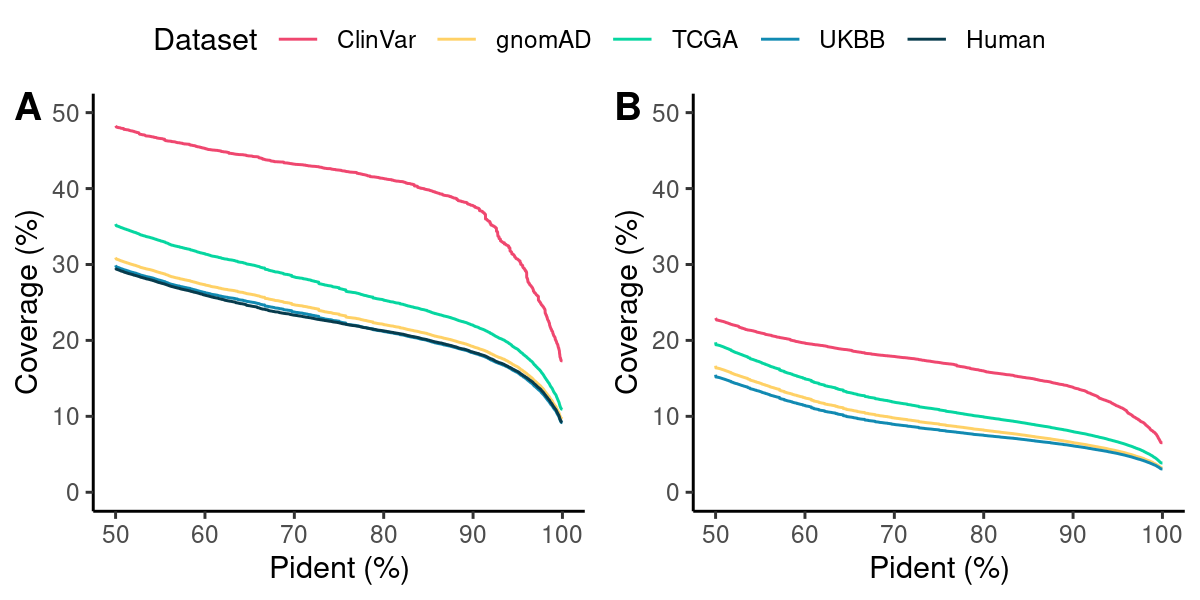

**Figure S6. Analysis of the relationship between percent identity (Pident) and the structural coverage of mapped missense variants.** (A) Structural coverage and (B) protein interface coverage are shown for each of the four studies (ClinVar, gnomAD, TCGA, and UKBB, color-coded), as well as for all residues in the human proteome (in black). Both plots consider results only for the proteins that were included in the mapping step (i.e.: 19,263).

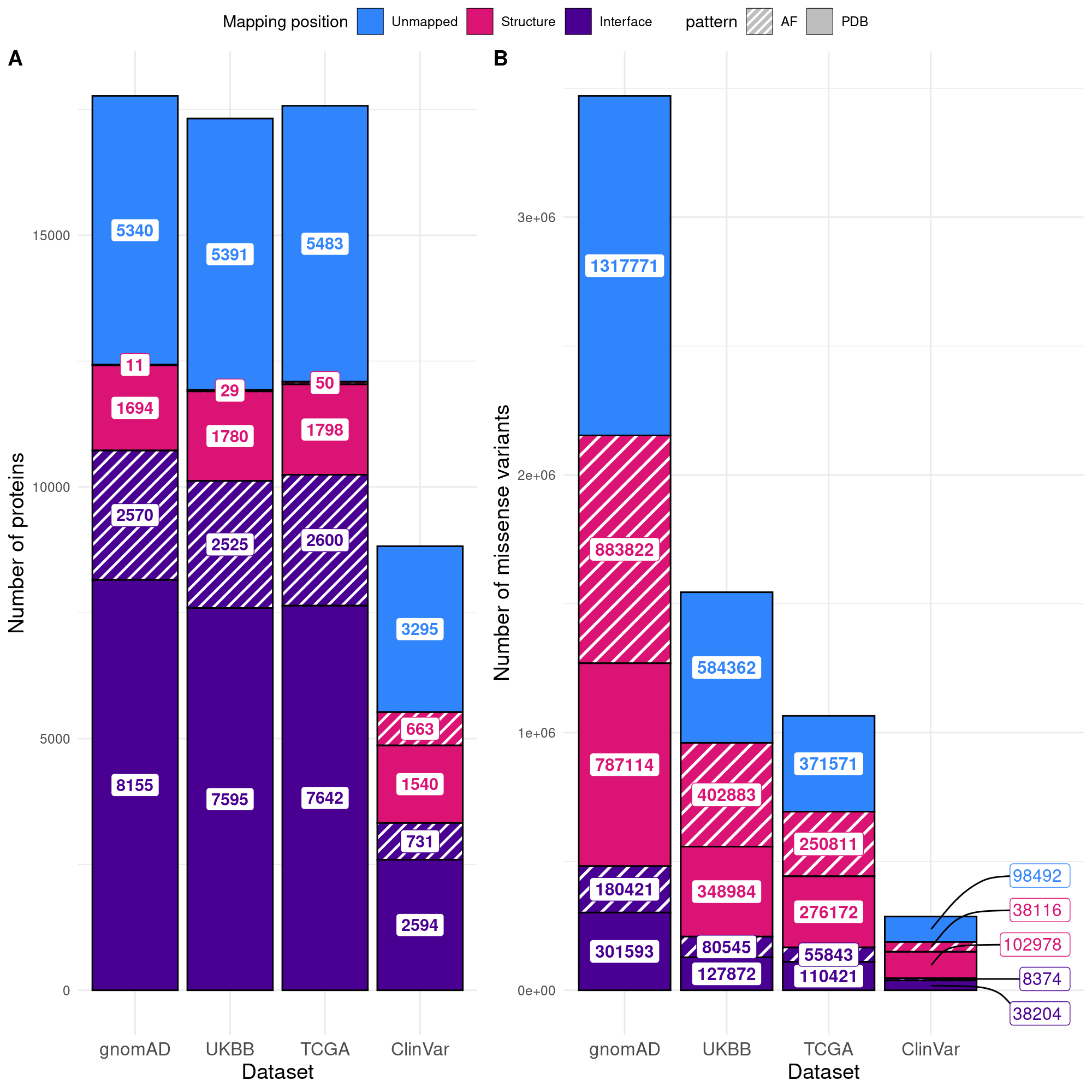

**Figure S7. Comparison of missense mutation structural coverage across principal protein isoforms of the human genome in four studies (ClinVar, UKBB, TCGA, and gnomAD). (A)** Barplot illustrating the number of proteins with at least one mapped missense variant in either the interface or the rest of the structure. **(B)** Distribution of structural coverage for missense variants across the human proteome. In both figures, stacked bars with patterns represent the additional structural coverage provided by AlphaFold models (see Methods). The "Unmapped" category indicates variants for which no structural coverage was identified (see Methods).

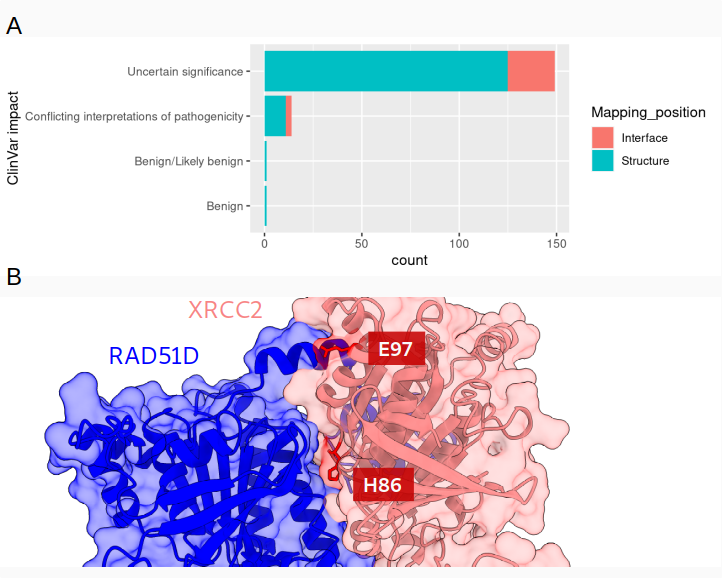

**Figure S8. Integration of AlphaFold models reveals interpretation of variants on mapping to interfaces. (A)** ClinVar impact predictions for mutations mapped to the AlphaFold model of the hu.MAP XRCC2 (O43543)-RAD51D (O75771) protein complex (pDockQ= 0.696, interface pLDDT = 86.85). While most are classified as uncertain significance with conflicting interpretations of pathogenicity, some can now be assessed by examining their interface impact in vitro or in silico. **(B)** Structure of the AlphaFold model of the hu.MAP XRCC2-RAD51D protein complex (pink and blue, respectively). XRCC2 mutations H86N and E97Q (highlighted in red), categorized as "probably damaging" in TCGA, are located in the protein complex interface.

**References**

1.	Via, A., Zanzoni, A. & Helmer-Citterich, M. Seq2Struct: a resource for establishing sequence-structure links. *Bioinforma. Oxf. Engl.* **21**, 551–553 (2005).

2.	Martin, A. C. R. Mapping PDB chains to UniProtKB entries. *Bioinformatics* **21**, 4297–4301 (2005).

3.	Wernersson, R., Rapacki, K., Stærfeldt, H.-H., Sackett, P. W. & Mølgaard, A. FeatureMap3D—a tool to map protein features and sequence conservation onto homologous structures in the PDB. *Nucleic Acids Res.* **34**, W84–W88 (2006).

4.	David, F. P. A. & Yip, Y. L. SSMap: a new UniProt-PDB mapping resource for the curation of structural-related information in the UniProt/Swiss-Prot Knowledgebase. *BMC Bioinformatics* **9**, 391 (2008).

5.	Stank, A., Richter, S. & Wade, R. C. ProSAT+: visualizing sequence annotations on 3D structure. *Protein Eng. Des. Sel.* **29**, 281–284 (2016).

6.	Wang, J. *et al.* G2S: a web-service for annotating genomic variants on 3D protein structures. *Bioinformatics* **34**, 1949–1950 (2018).

7.	Faezov, B. & Jr, R. L. D. PDBrenum: A webserver and program providing Protein Data Bank files renumbered according to their UniProt sequences. *PLOS ONE* **16**, e0253411 (2021).

8.	Velankar, S. *et al.* SIFTS: Structure Integration with Function, Taxonomy and Sequences resource. *Nucleic Acids Res.* **41**, D483-489 (2013).

9.	Ryan, M., Diekhans, M., Lien, S., Liu, Y. & Karchin, R. LS-SNP/PDB: annotated non-synonymous SNPs mapped to Protein Data Bank structures. *Bioinformatics* **25**, 1431–1432 (2009).

10.	Douville, C. *et al.* CRAVAT: cancer-related analysis of variants toolkit. *Bioinforma. Oxf. Engl.* **29**, 647–648 (2013).

11.	Wang, D. *et al.* SNP2Structure: A Public and Versatile Resource for Mapping and Three-Dimensional Modeling of Missense SNPs on Human Protein Structures. *Comput. Struct. Biotechnol. J.* **13**, 514–519 (2015).

12.	Solomon, O. *et al.* G23D: Online tool for mapping and visualization of genomic variants on 3D protein structures. *BMC Genomics* **17**, 681 (2016).

13.	Gress, A., Ramensky, V., Büch, J., Keller, A. & Kalinina, O. V. StructMAn: annotation of single-nucleotide polymorphisms in the structural context. *Nucleic Acids Res.* **44**, W463-468 (2016).

14.	Modeling Cancer Mutations in 3-D. *Cancer Discov.* **7**, 787–788 (2017).

15.	Konc, J., Skrlj, B., Erzen, N., Kunej, T. & Janezic, D. GenProBiS: web server for mapping of sequence variants to protein binding sites. *Nucleic Acids Res.* **45**, W253–W259 (2017).

16.	Radusky, L. *et al.* VarQ: A Tool for the Structural and Functional Analysis of Human Protein Variants. *Front. Genet.* **9**, (2018).

17.	Stephenson, J. D., Laskowski, R. A., Nightingale, A., Hurles, M. E. & Thornton, J. M. VarMap: a web tool for mapping genomic coordinates to protein sequence and structure and retrieving protein structural annotations. *Bioinformatics* **35**, 4854–4856 (2019).

18.	Laskowski, R. A., Stephenson, J. D., Sillitoe, I., Orengo, C. A. & Thornton, J. M. VarSite: Disease variants and protein structure. *Protein Sci. Publ. Protein Soc.* **29**, 111–119 (2020).

19.	Ferla, M. P., Pagnamenta, A. T., Koukouflis, L., Taylor, J. C. & Marsden, B. D. Venus: Elucidating the Impact of Amino Acid Variants on Protein Function Beyond Structure Destabilisation. *J. Mol. Biol.* **434**, 167567 (2022).

20.	Krebs, F. S. *et al.* Swiss-PO: a new tool to analyze the impact of mutations on protein three-dimensional structures for precision oncology. *Npj Precis. Oncol.* **5**, 1–9 (2021).

21.	Mooney, S. D. & Altman, R. B. MutDB: annotating human variation with functionally relevant data. *Bioinformatics* **19**, 1858–1860 (2003).

22.	Luu, T.-D. *et al.* MSV3d: database of human MisSense Variants mapped to 3D protein structure. *Database J. Biol. Databases Curation* **2012**, bas018 (2012).

23.	Cerami, E. *et al.* The cBio cancer genomics portal: an open platform for exploring multidimensional cancer genomics data. *Cancer Discov.* **2**, 401–404 (2012).

24.	Mosca, R. *et al.* dSysMap: exploring the edgetic role of disease mutations. *Nat. Methods* **12**, 167–168 (2015).

25.	Porta-Pardo, E., Hrabe, T. & Godzik, A. Cancer3D: understanding cancer mutations through protein structures. *Nucleic Acids Res.* **43**, D968-973 (2015).

26.	Woodard, J., Zhang, C. & Zhang, Y. ADDRESS: A Database of Disease-associated Human Variants Incorporating Protein Structure and Folding Stabilities. *J. Mol. Biol.* **433**, 166840 (2021).

27.	Tang, Z.-Z. *et al.* PSCAN: Spatial scan tests guided by protein structures improve complex disease gene discovery and signal variant detection. *Genome Biol.* **21**, 217 (2020).

28.	Niknafs, N. *et al.* MuPIT interactive: webserver for mapping variant positions to annotated, interactive 3D structures. *Hum. Genet.* **132**, 1235–1243 (2013).

29.	Laddach, A., Ng, J. C. F. & Fraternali, F. Pathogenic missense protein variants affect different functional pathways and proteomic features than healthy population variants. *PLOS Biol.* **19**, e3001207 (2021).

30.	Waterhouse, A. M., Procter, J. B., Martin, D. M. A., Clamp, M. & Barton, G. J. Jalview Version 2--a multiple sequence alignment editor and analysis workbench. *Bioinforma. Oxf. Engl.* **25**, 1189–1191 (2009).
