## Supplementary material for "3Dmapper: A Command Line Tool For BioBank-scale Mapping Of Variants To Protein Structures": 3D mapper - Tutorial

### 3Dmapper Documentation

### Table of contents


- Home
- Roadmap
- Installation
- Basic commands
  - makestructuraldb
  - makevariantsdb
  - mapper
  - makevisualization
- Tutorial mode
  - Step 1. Generation of a Protein Structural DB
  - Step 2. Generation of a Variant DB
  - Step 3. Map Variants with Structural Database
  - Step 4. Results Visualization
- Parallelization

### Home

Welcome to the 3Dmapper wiki!

#### Motivation

To understand biological processes, interpreting genomic data is crucial. Protein structures aid in this interpretation by providing functional context for genomic coding regions. Mapping genes to proteins is simple, but inconsistencies in data formats can make it a tedious and error-prone process. Over the last 20 years, more than 20 different tools or databases have been developed to automatically map annotated positions and variants to protein structures. However, most of these methods are web-based tools, which is not ideal for dealing with large-scale genomic data. To address this, we introduce 3Dmapper, a standalone Python/R command line tool that systematically maps annotated protein positions or variants to protein structures.

#### The tool

3Dmapper is a command-line tool that integrates structural and genomic data to identify and visualize variants on protein structures. The pipeline consists of four steps, including the generation of a structural database, variant annotation and splitting, variant mapping to protein structures, and visualization with ChimeraX. 3Dmapper enables users to identify the functional effects of variants, their locations in relation to the protein structure, and ultimately, their potential involvement in disease. The pipeline is customizable and can be used with any protein structure, facilitating the identification of variants in proteins of interest.

### Roadmap

The figure below provides a visual summary of the main steps involved in using 3Dmapper. Starting from the generation of the structural database and variant annotation files, the pipeline proceeds with the mapping of the variants onto the protein structures and ends with the visualization of the results using ChimeraX.

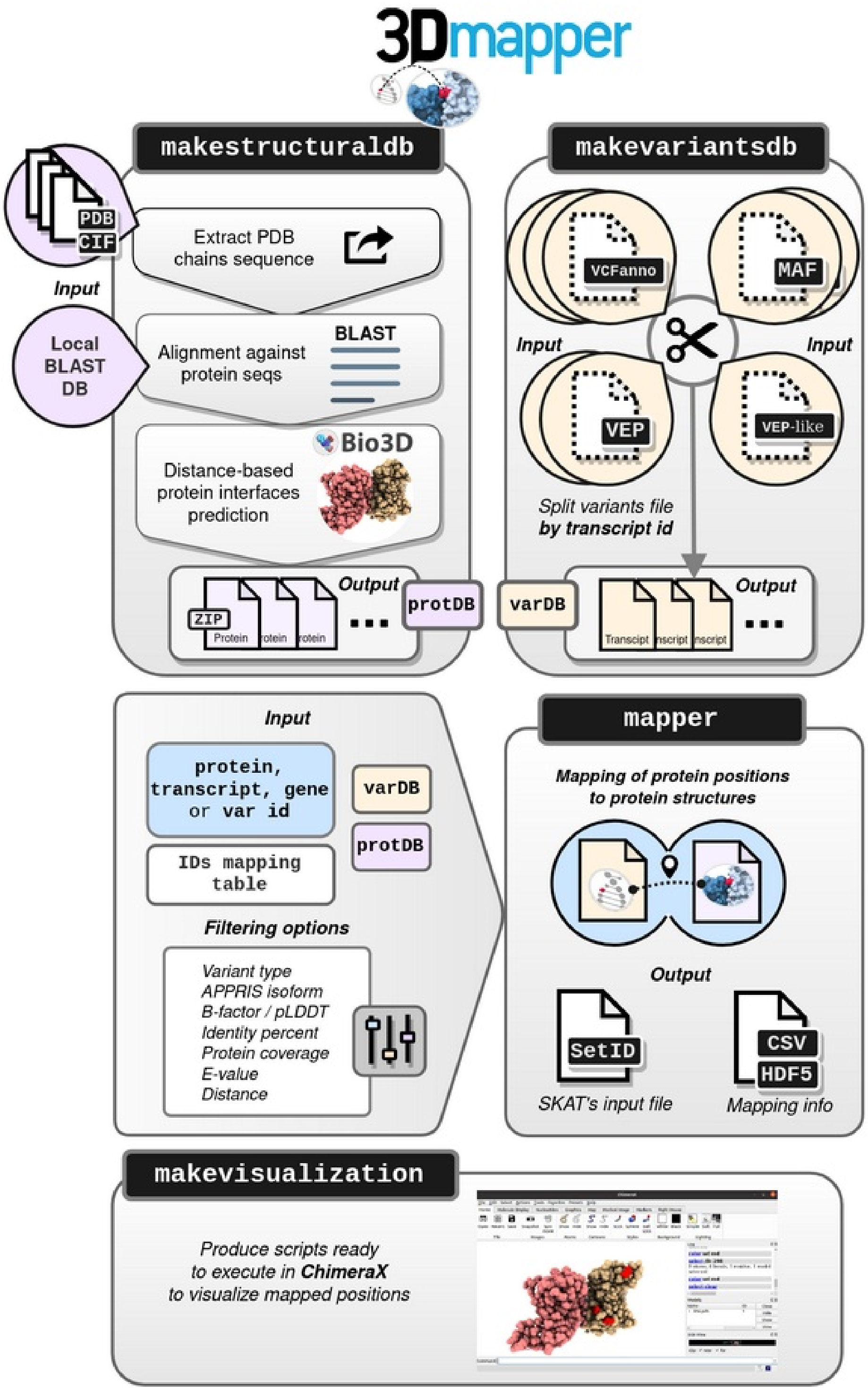

### Installation

#### Dependencies

##### Manual installation

Prior 3Dmapper installation, the following dependencies MUST be installed.

- BLAST standalone software version >= 2.6. Follow these instructions to download and use the command line tool.
- Python > 3.6
- R version > 3.5
- GNU parallel

#### 3Dmapper command line application installation

Execute in Terminal the following code:

```
git clone https://github.com/vicruiser/3Dmapper.git
cd 3Dmapper
pip install .
sh r_dependencies.sh
```

Congratulations, 3Dmapper is ready to use!

### makestructuraldb

To map variants or positions onto protein structures, we need a reference database of structural information. This database is generated using the `makestructuraldb` command.

#### What `makestructuraldb` does?

The `makestructuraldb` command performs several tasks:

1. **Extracts PDB chain sequences**: `makestructuraldb` extracts the amino acid sequences of each chain in the PDB files provided. This is necessary because we need to compare the sequences of the proteins in our dataset to those in the PDBs.
2. **BLAST PDB chain sequences against target proteome**: The chain sequences extracted from the PDBs are compared against the proteome of interest using BLAST. This step involves two parts:
   - **Generate BLAST database**: First, we need to create a BLAST database from the proteome of interest. This is done using the makeblastdb command, which accepts a FASTA file containing the protein sequences.
   - **BLAST search**: the BLAST search is done automatically using the PDB chain sequences as queries and the target proteome as the subject. The search is performed using the blastp program from the BLAST suite of tools.
3. **Filtering of BLAST hits**: After running the BLAST search, we filter the hits based on user-defined criteria such as e-value, identity percent, and coverage. This step ensures that only high-quality hits are used to create the reference database.
4. **Retrieves structural data of hits**: For the hits that pass the filtering step, we retrieve the corresponding structural data. This includes information about the atomic coordinates of the protein.
5. **Calculates interfaces**: Finally, `makestructuraldb` calculates the interfaces between proteins, ligands, and nucleic acids based on a distance cutoff.

#### Input files

- **PDB file(s)** or **a file with a list of PDBs** in **.pdb** or **.cif** formats. Also accepts compressed files in **.gz** format.
- A BLAST database that was generated using the `makeblastdb` BLAST command.

#### Input arguments

The program accepts the following input arguments:

- `--pdb` (required): the path to one or more PDB files.
- `--blast_db` (required): the path to the proteome files, which are the output of makeblastdb.
- `-o/--out`: the output directory where results will be saved. By default, the directory is set to "./structural_db".
- `-d/--dist`: the inter-residue distance threshold in angstroms. The default value is 5.
- `-t/--int-type`: the interface definition. The options are: 'noh' (by default) to calculate distance considering heavy atoms only; 'calpha' to compute CA-CA distance; or 'cbeta' to measure distances between CB-CB (CA in the case of Glycine).
- `-e/--evalue`: the e-value threshold in BLAST. The default value is 1e-7.
- `--pident`: the percent identity threshold between the query (PDB chain) and hit (protein) sequences. The default value is 20 percent.
- `-c/--coverage`: the percent coverage threshold of the protein sequence (how much of the protein sequence is covered by the PDB sequence). The default value is 0 percent.
- `--interaction`: the interaction type. The options are: 'protein', 'ligand', 'nucleic', a combination of the previous separated by space, or 'all' (by default).
- `--biolip`: consider the BioLiP list (https://zhanggroup.org/BioLiP/ligand_list) to remove artifact ligands? The default value is False.
- `-p/--parallel`: parallelize the process. The default value is False.
- `-j/--jobs`: the number of jobs to run in parallel. The default value is 1.

### Output

The output is a set of compressed files, each containing the structural data for individual proteins that obtained hits in the filtering step. These files will be used in subsequent steps of the workflow to map variants or positions onto the corresponding protein structures.

In summary, the following directories and files are generated:

```
|__structural_db
      |______makestructuraldb.report
      |______pdb_chainseqs
      |        |______PDB_prot_nuc_lig_db.txt
      |______structuralDB
               |______prot_ID1.txt.gz
               |______prot_ID2.txt.gz
               |______...
               |______prot_IDn.txt.gz
```

The program creates a default directory named "structural_db" that contains two files: "makepsdb.report", which provides details about the command executed, and the "structuralDB" directory, which stores the protein structural database categorized by protein ID. Another file generated by the program is "PDB_prot_nuc_lig_db.txt", which contains information about the chains present in the processed PDBs. This information includes their names, lengths, and count.

Each of the files in the structurealDB directory consist of a 26 column tab-delimited file. In all cases, **"PDB chain"** refers to the extracted PDB chain or query sequence from each PDB file and **"Protein"** refers to the hit sequence found with the BLAST search against the target proteome. A more detailed description of the meaning of each column ID is specified in the table below.

| Column name | Notes |
|---|---|
| **Protein_accession** | Target protein ID Length of the target protein sequence |
| **Protein_length** | Length of protein sequence |
| **Protein_position** | Positions relative to the target protein sequence |
| **Protein_aa** | Amino acids corresponding to the target protein positions |
| **PDB_code** | PDB ID |
| **PDB_chain** | ID of the template PDB protein chain |
| **PDB_chain_length** | Length of PDB chain sequence |
| **PDB_3D_position** | Position in the PDB chain **structure** |
| **PDB_seq_position** | Position in the PDB chain **sequence** |
| **PDB_aa** | Amino acids corresponding to both the PDB sequence and 3D positions |
| **Evalue** | E-value of the alignment between the query or PDB chain sequence and the target protein |
| **Pident** | Identity percent between the query (PDB chain) and the target sequence (protein). |
| **Protein_coverage** | Coverage (%) of the target protein by the PDB chain sequence |
| **Length_alignment** | Total length of the alignment between the query or PDB chain sequence and the target protein |
| **Interaction_type** | Type of interface interaction: "protein","nucleic" or "ligand". NA means no interaction which represents the positions of the rest of the structure |
| **PDB_interacting_chain** | Interacting PDB chain ID with the template PDB chain. NA means no interaction which represents the positions of the rest of the structure |
| **PDB_interacting_3D_position** | Position in the interacting PDB chain **structure** |
| **PDB_interacting_aa** | Amino acids corresponding to the interacting PDB structure positions |
| **Interface_min_distance** | Minimum existing distance between the pair of selected positions participating in the interface |
| **PDB_B_factor** | Minimum B factor (or pLDDT in the case of AF2 models) observed in each PDB 3D position |

| Column name | Notes |
| --- | --- |
| **PDB_interacting_B_factor** | Minimum B factor (or pLDDT in the case of AF2 models) observed in each PDB interacting 3D position |
| **Protein_alignment_start** | Alignment start position in target protein sequence |
| **Protein_alignment_end** | Alignment end position in target protein sequence |
| **PDB_alignment_start** | Alignment start position in PDB chain protein sequence |
| **PDB_alignment_end** | Alignment end position in PDB chain protein sequence |
| **Structure_feature_id** | As `PDB_code` _ `Protein_accession` _ `PDB_chain` _ `PDB_interacting_chain` _ `Interaction_type` |

### makevariantsdb

To accelerate the mapping process, we need to split variants or annotated positions files by individual transcript IDs.

#### What `makevariantsdb` does?

Splits the variants/positions file by transcript ID into individual files using `awk` and `sed` command lines.

#### Input files

- **Variants file**: The input annotated genomic variants file must be either in VCF, VEP or MAF default format. Additionally, a VEP-like format is admissible which must contain at least the following columns:
  - Uploaded_variation
  - Gene
  - Feature
  - Consequence
  - Protein_position
  - Amino_acids
- **Positions file**: If we want to *map a set of positions to protein structures that are NOT mutations*, we have to create a "positions" file. Its format imitates the one of variants files. More specifically, the easiest way to generate a "positions file" is to replicate the VEP-like format, keeping the same column names but adding some modifications.

An example of a position file would be:

- Uploaded_variation: contains the position ID
- Gene: can be aprotein ID
- Feature: can be a protein ID
- Consequence: description of this position, if any. Examples could be "conserved_position", "functional_position", etc
- Protein_position: position in the protein sequence
- Amino_acids: corresponding amino acid

This file it is not only useful to map positions or variants to protein structures, but also to find analogous positions of one protein in other protein structures across evolution. For example, if we have identified interesting positions in a protein, we can map this positions not only to the true structure, but also to other protein homologs, avoiding the obstacle of manual alignment and mapping.

#### Input arguments

- `--varfile, -vf`: input VCF, VEP or VEP-like file(s).
- `--maf_file, -maf`: input MAF file(s).
- `--force, -f`: force to overwrite? Inactive by default.
- `--out, -o`: output directory. Default is the current directory.
- `--sort, -s`: sort input file to split. It speeds up the splitting step.
- `--parallel, -p`: speed up running time. Depends on GNU Parallel. O. Tange(2011): GNU Parallel - The Command-Line Power Tool, login: The USENIX Magazine, February 2011: 42-47.
- `--jobs, -j`: number of jobs to run in parallel. Default is 1.

#### Output

The following directories and files are generated:

```
|__DBs
     |_____makevariantsdb.log
     |_____makevariantsdb.report
     |_____varDB
            |_____transcript_ID1.txt
            |_____transcript_ID2.txt
            |_____...
            |_____transcript_IDn.txt
```

By default, a directory called *DBs* is created containing files *makevariantsdb.log* and *makevaraintsdb.report*, which inform about the the executed command and the *varDB* directory which contains the splitted files by individual transcript ID.

### mapper

#### What `mapper` does?

It merges data of protein variants to related protein structures. Specifically, it generates mapping between IDs of choice, such as protein, transcript, gene or variant IDs, using a conversion file provided by the user.

#### Input data

The `mapper` command line tool requires specific input data to generate the mapping of protein, transcript, gene or variant IDs. Here is a breakdown of the input data required:

- **Interfaces database directory path**: This is a directory path to the interfaces database that was previously generated using the makestructuralsdb command line tool.
- **Variants database directory path**: This is a directory path to the variants database that was previously generated using the makevariantsdb command line tool.
- **IDs of choice**: This can be protein, transcript, gene, or variant IDs that the user wants to generate the mapping for.
- **ID mapping file**: A CSV file that contains the conversion of protein, transcripts and gene IDs, and optionally APPRIS isoforms IDs. This file is manually generated by the user based on their requirements. The file must have three columns with fixed names - "geneID", "transcriptID", and "protID". It should contain the conversion between these 3 IDs, where the IDs appearing in the structural dataset and the positions/variants files should be present. For instance, if the structural dataset was generated with a UniProt target proteome, then the protID column would contain UniProt IDs. If a variants file generated with VEP was used, then the geneID and transcriptID would correspond to Ensembl IDs.

It is essential to provide accurate input data to obtain the desired output from the `mapper` command line tool.

#### Input arguments

Input arguments of the program:

- `-pid, --prot-id`: one or more IDs of protein, transcripts or genes provided via command line or from a file
- `-vid, --var-id`: single or list of variants ids provided via command line or from a file
- `-psdb`: interfaces database directory (required)
- `-vdb`, --vardb: variants database directory (required)
- `-o, --out`: output directory (default: ./3dmapper_results)
- `--id_mapping`: file that contains the conversion of protein, transcripts and gene IDs and optionally APPRIS isoforms IDs (required)
- `-i, --isoform`: if available in the ID mapping file, this parameter can filter by a single or a list of APPRIS isoforms. The principal isoform is set by default. Options are: principal1, principal2, ...
- `-c, --consequence`: filter by variant or position consequence type (default: None)
- `-d, --dist`: threshold of interface maximum distance allowed in angstroms. By default, the maximum value will be the one selected in makeinterfacedb
- `--pident`: threshold of sequence identity (percentage) (default: 20)
- `-e, --evalue`: threshold of evalue (default: None)
- `-f, --force`: force to overwrite? Inactive by default (default: False)
- `-a, --append`: two or more calls to the program write are able to append results to the same output file (default: False)
- `-p, --parallel`: parallelize process (default: False)
- `-j, --jobs`: number of jobs to run in parallel (default: 1)
- `-v, --verbose`: print progress (default: False)

- `-l, --location`: map all variants and detect their location (default: False)
- `-csv`: write the mapped data to a CSV file (default: False)
- `-hdf`: write the mapped data to an HDF5 file using HDFStore (default: False)

#### Output

The output of the mapper command depends on the selected format:

- CSV: Generates 4 files, containing the following mapping results:
    - Interfaces: Contains the mapping results for protein interfaces.
    - Structures: Contains the mapping results for protein structures.
    - Unmapped: Contains the unmapped variants.
    - Non-coding: Contains the non-coding variants.

This output option is useful for having all the results in the same file. However, it can become difficult to handle in downstream analysis when mapping many variants or using many structures due to the large file size.

- HDF5: Generates 4 directories, each containing individual files per protein, transcript, gene, or variant ID. The directories and their contents are as follows:
    - Interface: Contains individual mapping results for protein interfaces.
    - Structure: Contains individual mapping results for protein structures.
    - Unmapped: Contains individual unmapped variants.
    - Non-coding: Contains the non-coding variants.

This output option is useful for large-scale mapping as it eases downstream analysis. HDF5 files can be quickly read using the vaex library in Python.

### makevisualization

#### What `makevisualization` does?

It generates ChimeraX scripts (.cxc) based on a template script. Once you have the mapped positions or variants in the CSV files from the mapper command output, you can visualize your results in 3D using ChimeraX.

#### Input Data

- PDB Code ID(s) or PDB file(s).
- Interface and/or Structure mapping results generated by `mapper` in CSV format.

#### Input Arguments:

- `-p or --pdb_code` : PDB code/s to be found within the mapped file. If assembly is not included it will generate a script for all mapped assemblies of that PDB code.
- `--pdb_list` : Specifies that PDBs provided are in a file, separated by spaces, tabs, or new lines.
- `-i or --interface_positions` : File containing the variants mapped to interfaces generated by 3Dmapper.
- `-s or --structure_positions` : File containing the variants mapped to protein structure generated by 3Dmapper.
- `-o or --output` : Output folder in which the ChimeraX script/s will be saved.
- `-n or --name` : Base name for the ChimeraX scripts.
- `-it or --inter_type` : If interfaces are available, filter by specified interaction type (ligand, protein or nucleic).
- `-l or --lighting` : Select lighting option: full, soft or simple.
- `-bg or --background` : Select background option: white or black.
- `-ns or --no_silhouette` : Display will not present silhouettes.
- `-mol or --mol_style` : Select option for molecule style: ball, sphere, or stick.
- `-is or --itf_style` : Select option for molecule style of the interfaces displayed: ball, sphere, or stick.
- `-f or --force` : Force to overwrite?

#### Output

The program generates ChimeraX scripts (.cxc) that highlight the mapped interface variants in red and the rest of the structure in blue. It also colors the PDB chains per chain ID. One script is generated per selected PDB, allowing the user to focus on specific structures of interest. This feature helps to visualize the location of the mapped variants within the PDB structure and provides a better understanding of the impact of the variants on the protein function.

### Tutorial mode

**Welcome to the 3Dmapper tutorial!**

In the following steps, we will guide you through the process of using 3Dmapper to map variants or protein positions to the structure of proteins. This tool is incredibly useful for researchers who want to gain insights into the potential effects of genetic variants on protein structure and function.

In this tutorial, we will cover both the general use of 3Dmapper, as well as a specific case using the sample data provided in the example folder. Specifically, we will be working with variants in transcripts **ENST00000367182** and **ENST00000374005**, which will be mapped to their corresponding PDB structures. These structures include both experimental and AlphaFold2 PDBs, providing a comprehensive reference dataset for analysis.

All the necesary input data to run this tutorial and the corresponding output files can be found in the 3Dmapper Github repository.

Let's get started!

### Step 1. Generation of a Protein Structural DB

In the first part of the tutorial, we will generate a protein structural database related to two human transcripts with Ensembl IDs **ENST00000367182** and **ENST00000374005**.

#### Build a BLAST Protein Database

To build a BLAST protein database, follow these steps: 1) Retrieve the human proteome in FASTA format. You can use a public repository such as UniProt or Ensembl.

2) Generate the BLAST protein database using the following command in the terminal:

```
makeblastdb -in UP000005640_9606.fasta.gz -dbtype prot -out human_proteome_uniprot
```

This command will generate three files with the name "human_proteome_uniprot" and extensions .phr, .pin, and .psq. You can find them in the human_proteome_blastdb folder.

#### Download PDB files

If you want to retrieve structural data for proteins that currently do not have structures in PDB, you can download all the files in PDB and rely on sequence homology to find structural homologs. To reproduce our example, we downloaded PDBs and AlphaFold2 models related to the transcripts of interest. You can find these files in the pdbs folder.

#### Execute `makestructuraldb`

To execute makestructuraldb, follow these steps:

1. In Terminal, cd to the 1-makestructuraldb directory.
2. Execute the following command:

```
makestructuraldb --pdb input_pdbs.txt --blast_db human_proteome_blastdb/human_proteome_uniprot --pident 95
```

The inputs are a list of PDB files and the human_proteome BLAST database. Additionally, a threshold of 95% sequence percent identity was set for filtering BLAST hits to reduce the number of results.

#### Output

The corresponding output can be found in the structural_db folder. For the selected input PDBs, structural information was found for 11 proteins (see structuralDB).

### Step 2. Generation of a Variant DB

The second step of the pipeline involves generating a variants database and splitting the file by transcript ID. To do this, we use the command `makevariantsdb`. The input file variants.vep contains variants associated with the transcripts ENST00000367182 and ENST00000374005.

To generate variant databases follow these steps:

1. In Terminal, `cd to the 2-makevariantsdb directory.

2. Execute the following command to split the variants.vep file:

```
makevariantsdb -vf variants.vep
```

This will create files within the varDB directory. These files contain the variants associated with each transcript, respectively.

Verify that the files have been successfully generated by navigating to the DBs folder on Github. You should see the two files with names corresponding to the transcript IDs.

### Step 3. Map Variants with Structural Database

To map the variants with the previously generated structural database, we need to follow these steps:

1. Open the terminal and navigate to the 3-mapper directory in the 3Dmapper repository.

2. Execute the following command:

```
mapper -pid P09769 O15151 -psdb ../1-makestructuraldb/structural_db/structuralDB/ -vdb ../2-makevariantsdb/DBs/varDB/ --id_mapping d
ict_geneprot_GRCh38_uniprot.txt -csv -a -l
```

This command maps the variants to the protein structures in the structural database and produces four CSV files containing information about the variants based on whether they are mapped to the interface, structure, not mapped, or non-coding.

- Input files:
  - ID mapping talbe
  - structuralDB
  - varDB
- Once the command has finished executing, you can find the resulting CSV files in the 3-mapper directory. Note that the columns of these files contain information present in both the variants and the structural database files.

Congratulations! Mapping of variants to the structural database is now complete. Great job!

### Step 4. Results Visualization

To visualize the mapping results obtained in Step 3, execute the following code:

```
makevisualization -p list_pdbs.txt --pdb_list -i ../3-mapper/3dmapper_results/csv/InterfacePositions_pident20.0_isoform_all_consequence_all.csv --force -s ../3-mapper/3dmapper_results/csv/StructurePositions_pident20.0_isoform_all_consequence_all.csv -is sphere
```

The generated Chimera scripts are here. Below you can visualize examples of the execution of two of the scripts.

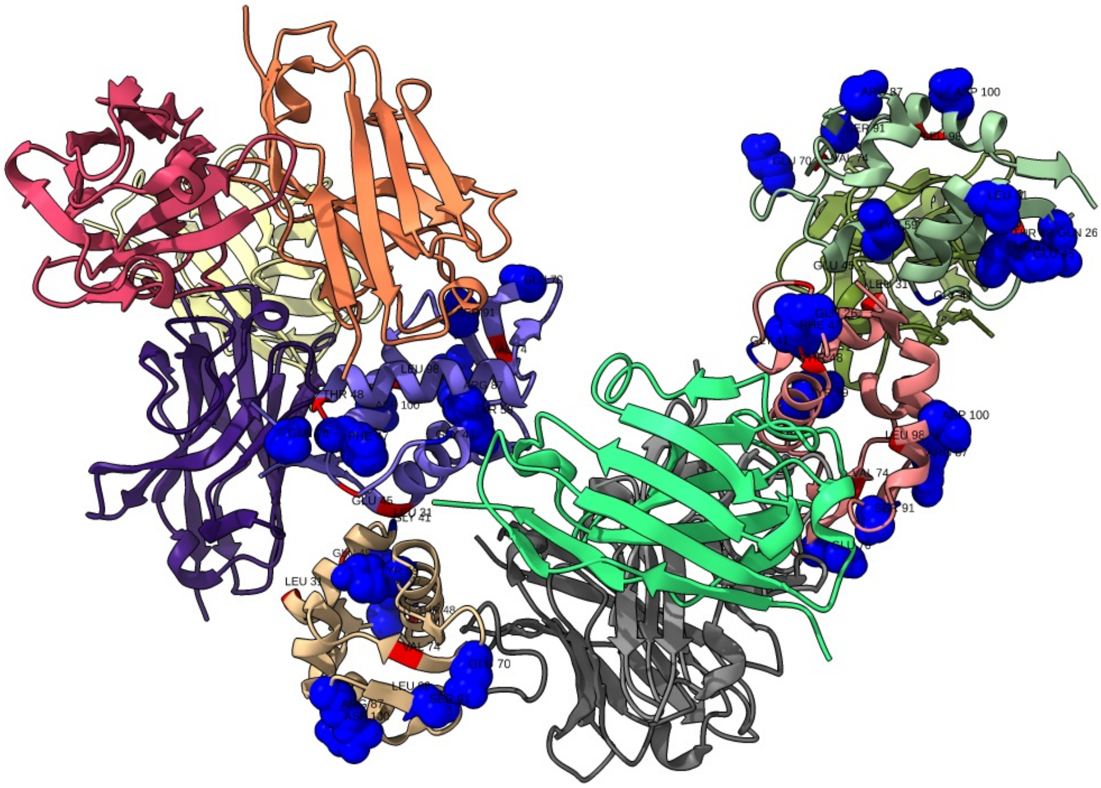


**Figure 1**. PDB 2vyr with variants mapped to the interface (in red) and to the structure (in blue).

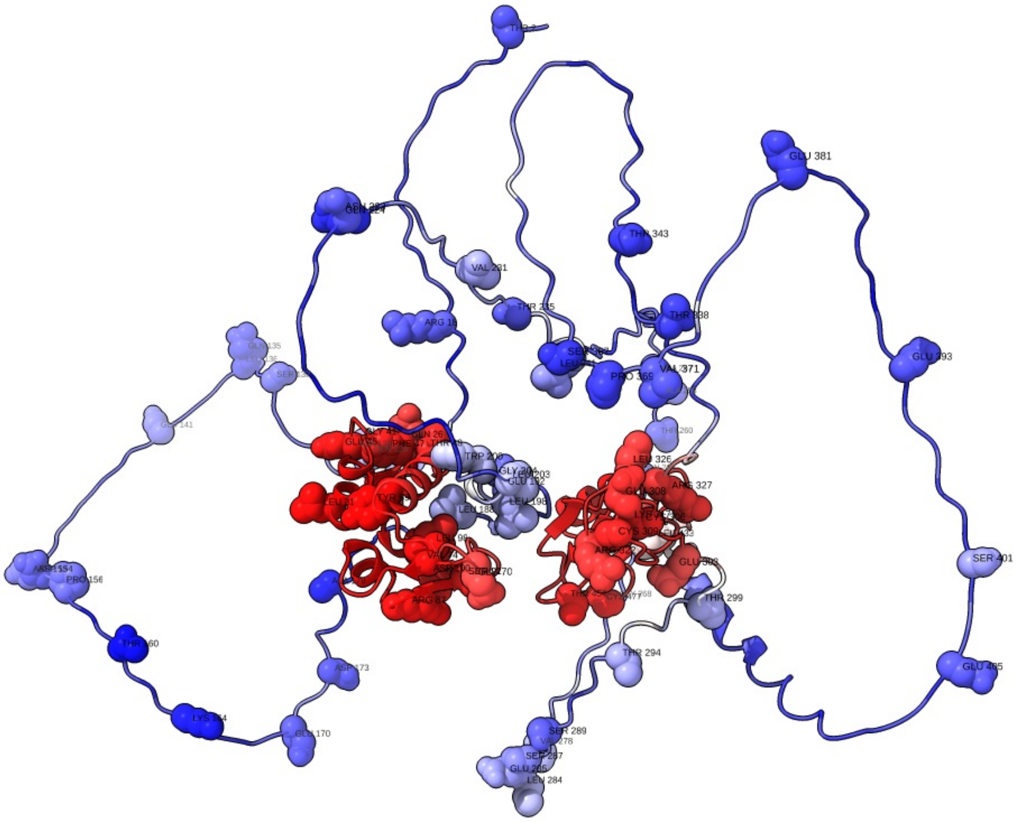


**Figure 2**. AlphaFold2 model O15151 with variants mapped to the structure (in blue) and chain colored in a red-blue gradient representing pLDDT scores (red, high pLDDT and blue low pLDDT).

### Parallelization

#### Parallelization with Joblib

To run `makestructuraldb` and `mapper` in parallel, activate the option `-p` or `--parallel` and specify the number of cores you want to use at the same time with `-j` or `--jobs`.

#### Parallelization in a Computer Cluster

In cluster computing, parallelization with Joblib may not be the optimal approach. Instead, you can submit individual tasks as a strategy of parallelization using job arrays or greasy, where the results are appended to the same output file. The option for parallelization with job arrays or greasy is available for `makestructuraldb` and `mapper` in 3Dmapper. Below are two examples of how to run this:

```
makestructuraldb -pdb file1.pdb --blast_db target_proteome_db
makestructuraldb -pdb file2.pdb --blast_db target_proteome_db
makestructuraldb -pdb file3.pdb --blast_db target_proteome_db
...
makestructuraldb -pdb fileN.pdb --blast_db target_proteome_db
```

```
mapper -pid protID1  -psdb structuralDB -vdb varDB -ids dict.txt  -csv -l
mapper -pid protID2  -psdb structuralDB -vdb varDB -ids dict.txt  -csv -l
mapper -pid protID3  -psdb structuralDB -vdb varDB -ids dict.txt  -csv -l
...
mapper -pid protIDN  -psdb structuralDB -vdb varDB -ids dict.txt  -csv -l
```

#### Parallelization with GNU parallel

The splitting process of `makevariantsdb` can be greatly improved by using GNU parallel, a powerful command-line tool for executing shell commands in parallel. You can activate this option by adding `-p` or `--parallel` to your command and specifying the number of cores you want to use at the same time with `-j` or `--jobs`.

To use GNU parallel with makevariantsdb, you need to manually install it first. Here's how you can install it on Ubuntu or Debian:

```
sudo apt-get update
sudo apt-get install parallel
```

Once you have installed GNU parallel, you can use it to split your input files into smaller chunks that can be processed in parallel by `makevariantsdb`. This can significantly speed up the overall process, especially for larger input files.

Here's an example of how to use GNU parallel with `makevariantsdb`:

```
makevariantsdb -vf variants.vep -p -j 4
```

In this example, the -p option activates parallel processing, and -j 4 specifies that we want to use 4 cores at the same time. You can adjust this number based on the available resources on your system.

Note that you need to have a multi-core CPU or access to a high-performance computing cluster to take advantage of parallel processing.
